# The DHODH Inhibitor PTC299 Arrests SARS-CoV-2 Replication and Suppresses Induction of Inflammatory Cytokines

**DOI:** 10.1101/2020.08.05.238394

**Authors:** Jeremy Luban, Rachel Sattler, Elke Mühlberger, Jason D. Graci, Liangxian Cao, Marla Weetall, Christopher Trotta, Joseph M. Colacino, Sina Bavari, Caterina Strambio-De-Castillia, Ellen L. Suder, Yetao Wang, Veronica Soloveva, Katherine Cintron-Lue, Nikolai A. Naryshkin, Mark Pykett, Ellen M. Welch, Kylie O’Keefe, Ronald Kong, Elizabeth Goodwin, Allan Jacobson, Slobodan Paessler, Stuart Peltz

**Author notes:** Authors contributed equally to this study (Jeremy Luban, Rachel Sattler, Elke Mühlberger). Edge BioInnovation Frederick, MD 21702. Pharmacology, Merck BMB11-110, 33 Avenue Louis Pasteur, Boston MA, 02115.

## Abstract

The coronavirus disease 2019 (COVID-19) pandemic has created an urgent need for therapeutics that inhibit the SARS-CoV-2 virus and suppress the fulminant inflammation characteristic of advanced illness. Here, we describe the anti-COVID-19 potential of PTC299, an orally available compound that is a potent inhibitor of dihydroorotate dehydrogenase (DHODH), the rate-limiting enzyme of the de novo pyrimidine biosynthesis pathway. In tissue culture, PTC299 manifests robust, dose-dependent, and DHODH-dependent inhibition of SARS CoV-2 replication (EC_50_ range, 2.0 to 31.6 nM) with a selectivity index >3,800. PTC299 also blocked replication of other RNA viruses, including Ebola virus. Consistent with known DHODH requirements for immunomodulatory cytokine production, PTC299 inhibited the production of interleukin (IL)-6, IL-17A (also called IL-17), IL-17F, and vascular endothelial growth factor (VEGF) in tissue culture models. The combination of anti-SARS-CoV-2 activity, cytokine inhibitory activity, and previously established favorable pharmacokinetic and human safety profiles render PTC299 a promising therapeutic for COVID-19.

## INTRODUCTION

The coronavirus disease 2019 (COVID-19) pandemic is a serious threat to public health. Severe acute respiratory syndrome coronavirus 2 (SARS-CoV-2), the causative agent of COVID-19, is a positive-sense, single-stranded-RNA virus of the *Coronaviridae* family that shares 79.5% sequence identity with SARS-CoV (Huang et al., 2020; Lu et al., 2020; Wu et al., 2020a; Wu et al., 2020b; Zhou et al., 2020b; Zhu et al., 2020). While both viruses are likely descendants of bat coronaviruses, the proximal source of this zoonotic virus is unknown (Zhou et al., 2020b). Since its first description at the end of 2019, SARS-CoV-2 has spread around the globe, infecting millions of people, and killing hundreds of thousands (https://coronavirus.jhu.edu). An urgent medical need exists for effective treatments for this disease.

In the early stages of COVID-19, the virus proliferates rapidly, and in some cases, triggers a cytokine storm - an excessive production of inflammatory cytokines (Quartuccio et al., 2020; Wiersinga et al., 2020). This uncontrolled inflammation can result in hyperpermeability of the vasculature, multi-organ failure, acute respiratory distress syndrome (ARDS), and death (Gupta et al., 2020; Jose and Manuel, 2020; Quartuccio et al., 2020; Wang et al., 2020a; Zhou et al., 2020a). Acute respiratory distress syndrome is one of the leading causes of mortality in COVID-19 (Moore and June, 2020). In addition, elevated levels of interleukin (IL)-6 and IL-17 are reported to be associated with severe pulmonary complications and death (Pacha et al., 2020; Ruan et al., 2020; Russell et al., 2020).

A therapeutic that can inhibit SARS-CoV-2 replication while attenuating the cytokine storm would be highly beneficial for both the early and late stages of COVID-19. Targeting the cellular de novo pyrimidine nucleotide biosynthesis pathway is one potential approach to treat both phases of COVID-19 as viral replication and over-production of a subset of inflammatory cytokines are controlled by pyrimidine nucleotide levels (Cheung et al., 2017; Xiong et al., 2020). Dihydroorotate dehydrogenase (DHODH), the rate-limiting enzyme in this pathway, is located on the inner membrane of mitochondria and catalyzes the dehydrogenation of dihydroorotate (DHO) to orotic acid, ultimately resulting in the production of uridine and cytidine triphosphates (UTP and CTP) (Figure 1) (Munier-Lehmann et al., 2013).

**Figure 1.**
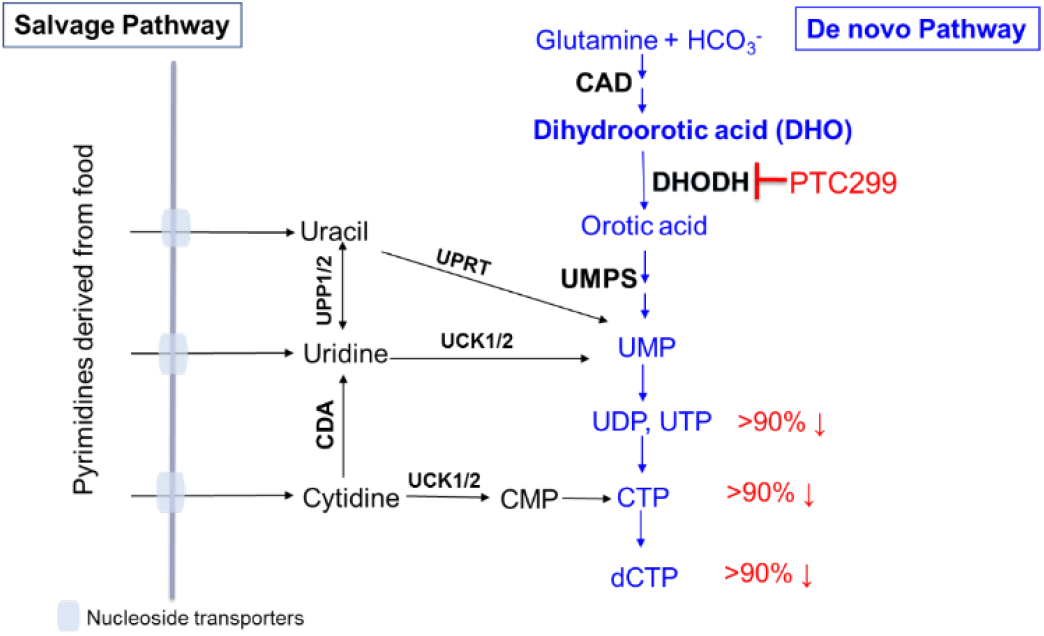
Pyrimidine Nucleotide Synthesis. Schematic of the salvage and de novo pathways of pyrimidine biosynthesis. The salvage pathway recycles pre-existing nucleotides from food or other extracellular sources and does not appear to be sufficient to support viral RNA replication. The percentages in red denote the respective extents to which the relative levels of specific nucleotides are reduced during PTC299 treatment (100 nM) of cultured HT1080 fibrosarcoma cells for 8 hours (Cao et al., 2019) Abbreviations: CAD, complex of the following three enzymes: carbamoyl-phosphate synthetase 2, aspartate carbamoyl transferase, and dihydroorotase; CDA, cytidine deaminase; CMP, cytidine monophosphate; CTP, cytidine triphosphate; DHO, dihydroorotate; UCK1/2, Uridine-cytidine kinase 1 and Uridine-cytidine kinase 2; UDP, uridine diphosphate; UMP, uridine monophosphate; UPP, uridine phosphorylase; UPRT, uracil phosphoribosyl transferase; UTP, uridine triphosphate.

The de novo pyrimidine biosynthesis pathway is also critical for the excessive production of a subset of inflammatory cytokines (including interferon-gamma, monocyte chemoattractant protein-1 [MCP-1], IL-5, and IL-6) in both cultured cells and animal models of viral infection (Cheung et al., 2017; Xiong et al., 2020). Consistent with DHODH being mechanistically central to COVID-19, a recent multi-omics study of drug targets for viral infections prioritized DHODH inhibition as one of the top three mechanisms to consider for SARS-CoV-2 treatment (Zheng et al., 2020).

PTC299 is an orally bioavailable potent inhibitor of DHODH (Cao et al., 2019). Treatment of cultured cells with PTC299 results in inhibition of DHODH activity, leading to increased levels of DHO, the substrate for the DHODH enzyme (Cao et al., 2019). Similar findings were documented in PTC299-treated cancer patients, where administration of PTC299 resulted in increased blood levels of DHO, indicating successful inhibition of DHODH in these patients (Cao et al., 2019). These results are consistent with increased DHO levels observed in Miller syndrome patients who carry mutations in the *DHODH* gene that reduce DHODH activity (Duley et al., 2016) and with studies showing that PTC299 not only inhibits vascular endothelial growth factor (VEGF) levels in cultured cells, but also normalized VEGF levels in cancer patients (Cao et al., 2019). VEGF is a stress-regulated cytokine the levels of which are increased in these patients (Plotkin et al., 2009).

In studies of over 300 human subjects, including healthy volunteers and oncology patients, PTC299 has manifested a favorable pharmacokinetic (PK) profile and has been generally well tolerated. The mechanism of action and the favorable PK and safety profiles of PTC299 support its investigation for use as a therapeutic for COVID-19. Here, we evaluated the in vitro antiviral activity of PTC299 against a panel of RNA viruses, with a specific focus on SARS-CoV-2, as well as the drug’s ability to attenuate the excessive production of inflammatory cytokines. Our results indicate that that PTC299 has considerable potential as a COVID-19 therapeutic.

## RESULTS

### PTC299 inhibits SARS-CoV-2 replication in vitro

The ability of PTC299 to inhibit SARS-CoV-2 replication was evaluated in vitro using two virus cell culture models. In the first model, Vero cells were preincubated for 24 hours with increasing concentrations of PTC299 and cells were subsequently infected with SARS-CoV-2 at a multiplicity of infection (MOI) of 0.05. Forty-eight hours after infection, viral protein was quantified by staining with antibody to the nucleocapsid protein of SARS-CoV-2 and with DAPI, which stains the cells nuclei, to determine cell number. The results demonstrated that PTC299 treatment led to a dose-dependent reduction in the levels of SARS-CoV-2 nucleocapsid protein (green) with an EC_50_ of 1.96 nM (Figures 2A and 2B). DAPI staining of cell nuclei (blue) showed that PTC299 did not reduce cell number (Figure 2A). These results indicate that PTC299 reduces viral load in the absence of cytotoxicity.

**Figure 2.**
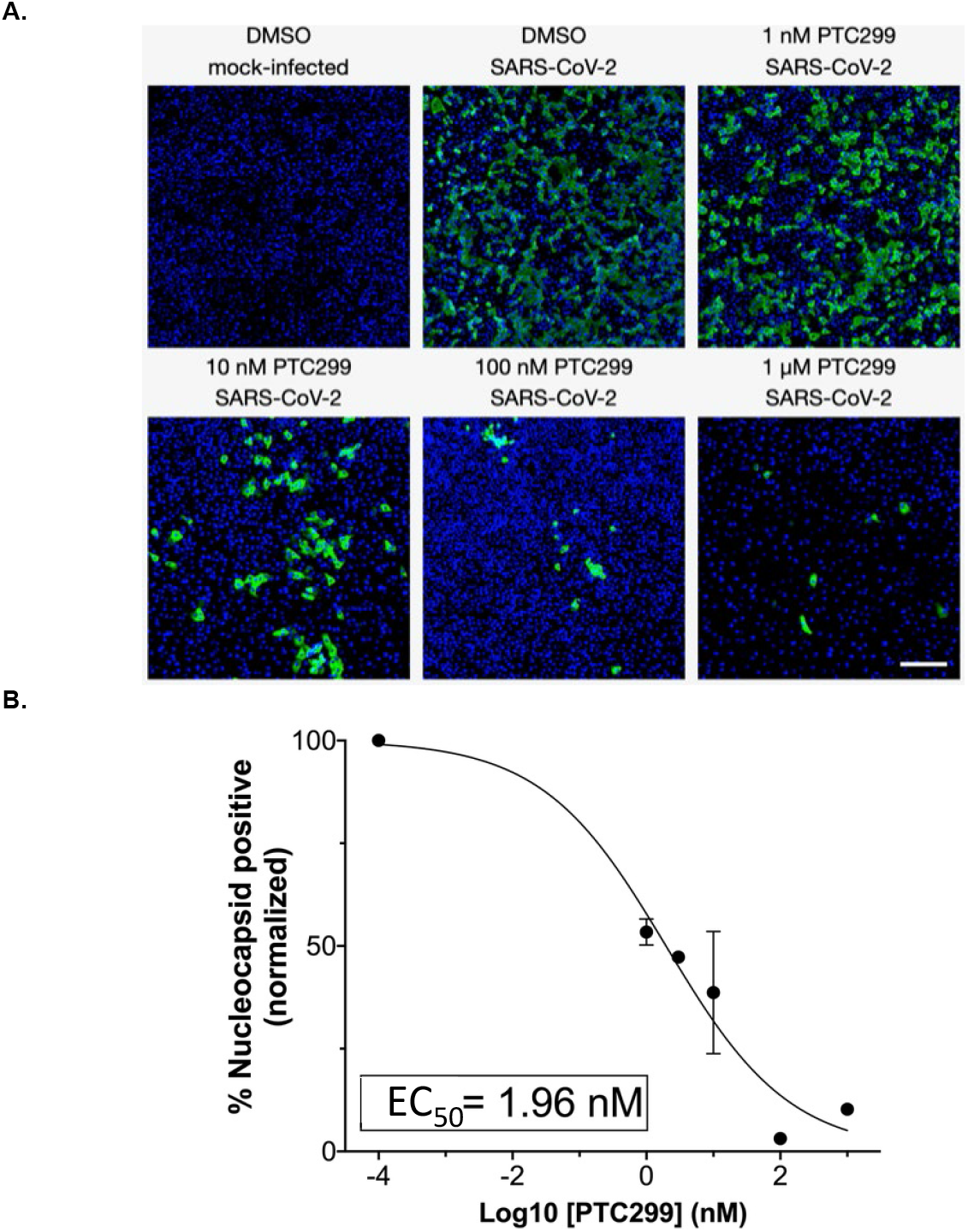
PTC299 inhibits SARS-CoV-2 replication. Legend: (A) Quantitative immunofluorescence analysis of SARS-CoV-2-infected Vero E6 cells treated with PTC299. PTC299 was added at concentrations ranging from 1 nM to 1 μM 30 minutes prior to infection of Vero E6 cells with SARS-CoV-2 (USA-WA1/2020) at a multiplicity of infection (MOI) of 0.1. At 48 hours post-infection, the cells were fixed, probed with antibodies against the nucleocapsid protein (NP) of SARS-CoV-2 and stained with AlexaFluor 488 conjugated secondary antibody. Nuclei were stained with DAPI. Images acquired in the Green (i.e., NP of SARS-CoV-2) and the Blue (DAPI) channels were overlaid and are displayed as indicated (note images corresponding to the 3 nM concentration of PTC299 were omitted for simplicity). Scale bar, 200 μm. (B) In order to quantify viral infection, images acquired in A were subjected to quantitative image analysis to determine the number of SARS-CoV-2 nucleocapsid positive cells at each concentration of PTC299 (i.e., 1, 3, 10, 100 nM and 1 μM). Normalized percent numbers of nucleocapsid positive cells were plotted against Log10 transforms of PTC299 concentrations and the IC_50_ was estimated using non-linear regression (goodness-of-fit R-squared = 0.9653). Data is plotted as the mean and standard deviation of 3 independent replicates.

In a second cell culture model, the efficacy of PTC299 in inhibiting SARS-CoV-2 replication was further evaluated. Vero cells were infected with SARS-CoV-2 at a MOI of 0.05 and PTC299 was added at increasing concentrations at various times from 18 hours pre-infection to 2 hours post-infection. Viral titer was quantified by collecting the medium from the infected cell cultures and determining the TCID_50_ (50% tissue culture infectious dose). The 50% and 90% effective concentrations (EC_50_ and EC_90_, respectively) of PTC299 were calculated. In parallel, the concentration of PTC299 necessary to reduce cell viability by 50% (cytotoxic concentration; CC_50_) in the absence of infection was determined by measuring intracellular ATP levels, and the selectivity index was calculated as CC_50_/EC_50_.

Preincubation of Vero cells with PTC299 at concentrations of 250, 500, and 1000 nM for 2 hours prior to viral infection reduced viral replication by approximately 3 orders of magnitude after 24 hours for all concentrations tested (Figure 3A). To determine whether the effect of PTC299 is a direct consequence of DHODH inhibition, excess uridine (100 μM) was added to the treatment media. Exogenously added uridine prevented the inhibition of SARS-CoV-2 replication by PTC299 (Figure 3B) consistent with the drug inhibiting DHODH to block viral replication. PTC299 also inhibited viral replication when added to the cells 18 hours prior to infection, added at the time of infection, or added 2 hours post-infection (see Supplemental Material Figure 1).

**Figure 3.**
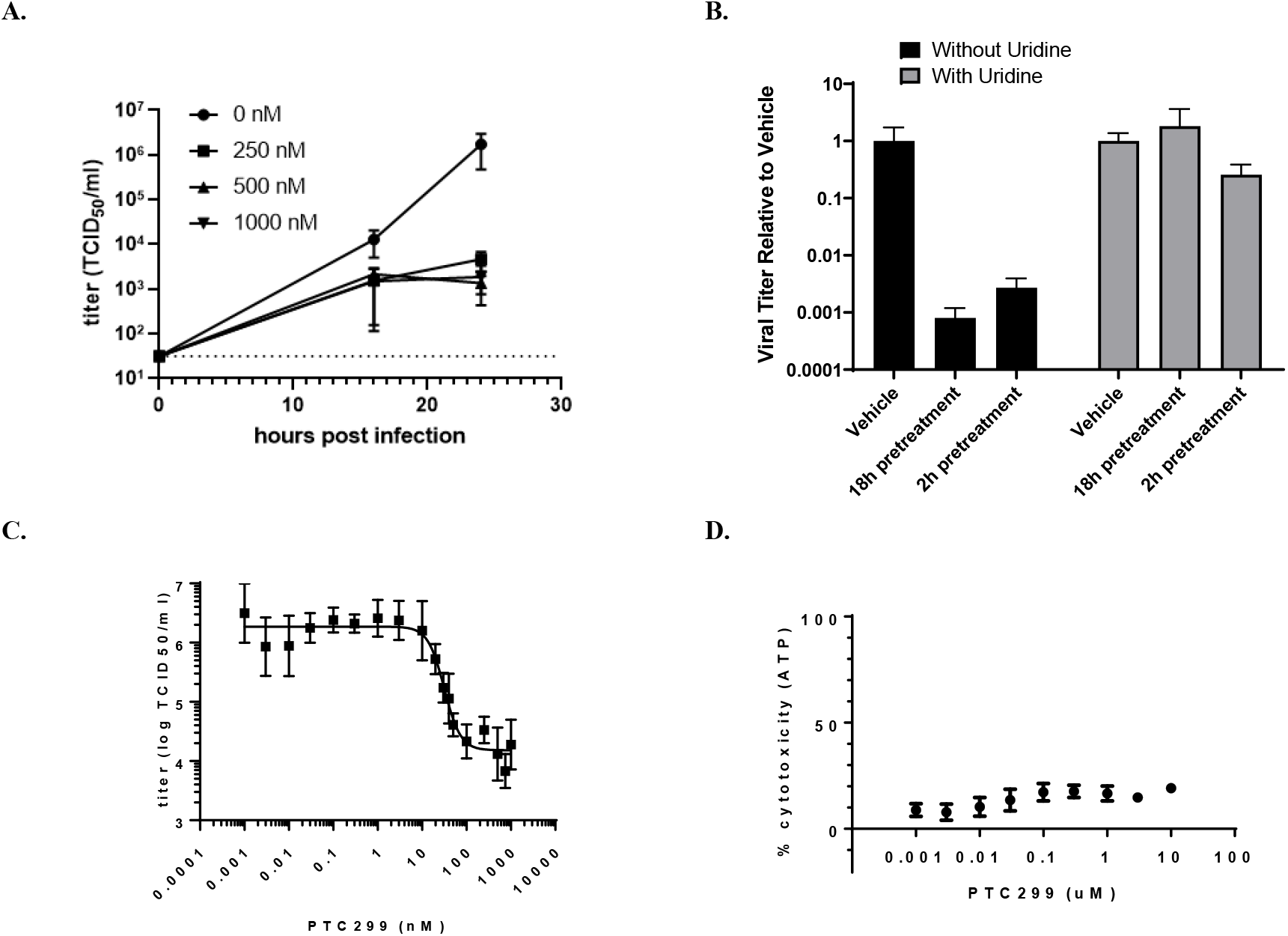
PTC299 decreases the titer of SARS-CoV-2 through the de novo pyrimidine synthesis pathway. Legend: (A) Quantification of the dose dependent inhibition of SARS-CoV-2 replication by PTC299. Vero cells were pretreated for 2 hours with the concentrations of PTC299 of 250, 500, or 1000 nM prior to the addition of SARS-CoV-2. Dashed line indicates the limit of quantification. (B). Uridine blocks the PTC299 inhibition of SARS-CoV2 replication. Vero cells were preincubated for 2 or 18 hours with PTC299 alone or with PTC299 plus 100 μM uridine before addition of SARS-CoV-2. (C) Determination of EC_50_ and EC_90_. Vero cells were preincubated with PTC299 at 0.003, 0.01, 0.10, 0.30, 1.0, 3.0, 10, 20, 30, 40, 50, 100, 250, 500, 750, and 1000 nM prior to addition of SARS-CoV-2. (D) PTC299 has little cytotoxic effect in Vero cells. Vero cells were preincubated for 2 hours with PTC299 at 0.001, 0.003, 0.010, 0.030, 0.10, 0.30, 1.0, 3.0, and 10 nM prior to addition of SARS-CoV-2. (A-D) Cells were inoculated with SARS-CoV-2 (USA-WA1/2020) at a MOI of 0.05. Viral titer was determined by collecting the medium from the infected cells and assessing the concentration required for 50% infectivity of Vero cells (the 50% tissue culture infectious dose assay or TCID_50_). Cytotoxicity was evaluated by measuring intracellular ATP (adenosine triphosphate) levels after 48 hours of culture. Data are plotted as the mean and standard deviation of 3 independent replicates.

The initial concentration range of PTC299 explored did not enable the determination of EC_50_ and EC_90_ for PTC299 against SARS-CoV-2. Therefore, a range of lower concentrations of PTC299 (from 1 pM to 1 μM) was evaluated. The EC_50_ and EC_90_ values were determined to be 2.6 nM and 53 nM, respectively (Figure 3C and Table 1), which is consistent with the findings of the immunofluorescence experiments described in Figure 2. The EC_50_, determined as the midpoint of the logarithmic curve, was 31.6 nM. Using a similar calculation based on the logarithmic curve, the EC_90_ was determined to be 80.3 nM. At the highest concentration of PTC299 tested (10 μM), the CC_50_ was determined to be >10 μM (Figure 3D and Table 1), indicating, similar to the immunofluorescence experiments described above, that PTC299 reduces viral load with little cell death.

**Table 1.**
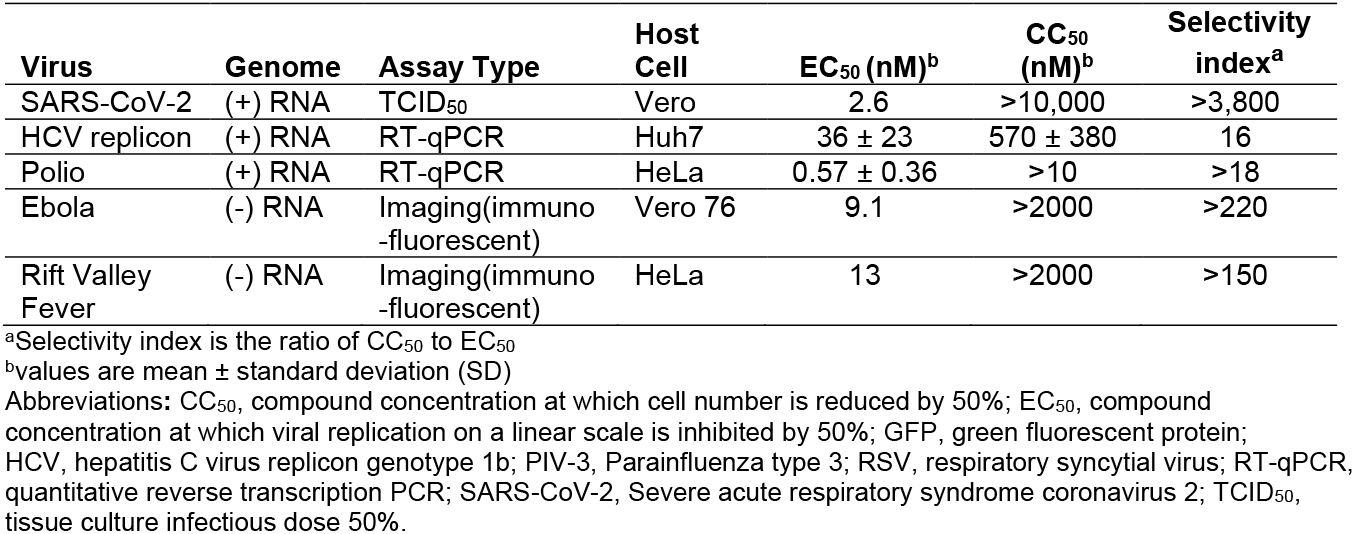
PTC299 has broad-spectrum antiviral activity.

### PTC299 has broad-spectrum antiviral activity

The results in Figures 2 and 3 indicate that PTC299 is a potent inhibitor of SARS-CoV-2 replication. Considering its mechanism of inhibiting cellular de novo pyrimidine synthesis and the established role of pyrimidine nucleotides in viral replication, we investigated the spectrum of activity of PTC299 against a panel of RNA viruses of both positive and negative genome polarity. PTC299 was found to have broad-spectrum antiviral activity against the RNA viruses tested (Table 1). PTC299 was a highly potent inhibitor of hepatitis C virus (HCV) replicon genotype 1b, poliovirus, Ebola virus, and Rift Valley fever virus (EC_50_ ≤36 nM). The CC_50_ values indicated minimal cytotoxicity in most of the viral infection models and were generally above the highest concentrations of PTC299 tested.

### PTC299 inhibits production of inflammatory cytokines

PTC299 was originally identified as a compound that modulates expression of stress-regulated proteins like VEGF and was only subsequently found to be a DHODH inhibitor (Cao et al., 2019; Weetall et al., 2016). DHODH inhibitors have been shown to suppress virally-induced inflammatory cytokine production (Xiong et al., 2020). The effect of PTC299 on cytokine production and release into the cell culture medium was assessed in the Biologically Multiplexed Activity Profiling (BioMAP) assay profiling platform. This system consists of co-cultures of peripheral blood mononuclear cells (PMBCs) with other cell types, including B cells, fibroblasts, and endothelial cells, allowing for the measurement of physiologically relevant biomarker readouts of the activity of a test compound (Berg et al., 2006; Kunkel et al., 2004a; Kunkel et al., 2004b). Each co-culture was stimulated to recapitulate relevant signaling networks that naturally occur in healthy human tissue and under certain disease states. For each system, cytotoxicity and cellular proliferation were assessed. PTC299 was added at increasing concentrations to each of the model systems one hour before cellular stimulation and remained present during the stimulation period for each co-culture. Cytokine production was subsequently assessed.

PTC299 was found to be a potent inhibitor of the production of a subset of immunomodulatory and inflammation-related cytokines that are associated with the stress response and poor prognosis in COVID-19 (Figure 4). In the SAg co-cell culture system, which models T-cell activation, PTC299 treatment resulted in significant decreases of MCP-1, CD40, and IL-8, as levels decreased after 24 hours of stimulation (Table 2). Compared with the control, 31% and 27% reductions in MCP-1 and IL-8, respectively, were observed when cells were incubated with 10 nM PTC299, and a 29% decrease in CD40 was detected with 100 nM PTC299 (all p values <0.01). A 23% reduction in endothelial cellular proliferation as measured by SRB (a measure of cell biomass) relative to control was seen following treatment with PTC299 at 100 nM (p<0.01) but not at 1 or 10 nM.

**Figure 4.**
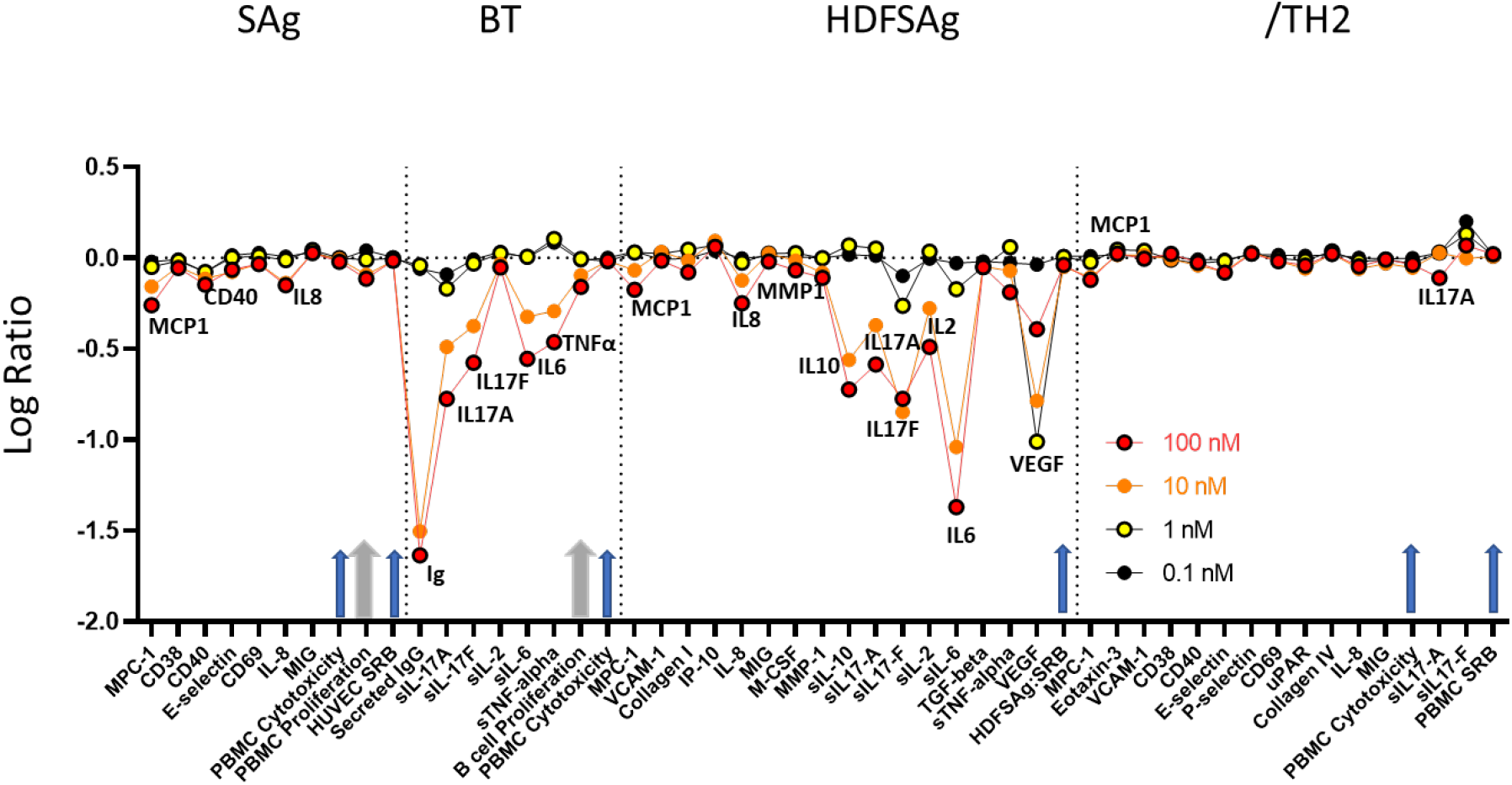
PTC299 Inhibits Production of Inflammatory Cytokines Measured Using the BioMAP Profile. Legend: The x-axis indicates the cytokines evaluated and the y-axis the log ratio of PTC299 treated vs. vehicle control. Arrows indicate evaluation of cell viability, cytotoxicity, and proliferation, with the two thicker grey arrows showing values that reached statistical significance at 100 nM PTC299. Biomarkers are annotated only when there was a significant difference between the effect of PTC299 and vehicle control (p<0.01), were outside of the significance envelope, and were >20% [log10 ratio>0.1]. The x-axis indicates the cytokines evaluated and the y-axis the log ratio of PTC299 treated vs. vehicle control. Arrows indicate evaluation of cell viability, cytotoxicity, and proliferation, with the two thicker grey arrows showing when values reached statistical significance at 100 nM PTC299. Abbreviations: BCR, B-cell receptor; BioMAP, Biologically Multiplexed Activity Profiling; BT System, co-culture of CD19+ B cells and PBMC that utilizes BCR stimulation and sub-mitogenic TCR stimulation; HDFSAg system, co-culture of human primary dermal fibroblasts and PBMC that is stimulated with sub-mitogenic TCR levels; Ig, immunoglobulin; IgG Immunoglobulin G; IL, interleukin; PBMC, Peripheral Blood Mononuclear Cell; SAg system, co-culture of endothelial cells and PBMC stimulated with mitogenic levels of TCR ligands; TCR, T-cell receptor; /TH2 system, co-culture of endothelial cells and Th2 blasts stimulated with TCR ligands and cytokines; Th2, T helper type 2; VEGF, vascular endothelial growth factor

**Table 2.** PTC299 Potently Inhibits Cytokines Production (p<0.01) – Summary of BioMap Profile Results.

| **Cytokine** | **Co-culture system** | **Dose (nM) with p<0.01** | **% Inhibition** |
| --- | --- | --- | --- |
| MCP-1 | SAG | 10, 100 | 31, 45 |
| CD40 | SAG | 100 | 29 |
| IL-8 | SAG | 10, 100 | 27, 29 |
| Endothelial Cell Proliferation | SAG | 100 | 23 |
| slgG | BT | 10, 100 | 97, 98 |
| sIL-17A | BT | 10, 100 | 68, 83 |
| s-IL-17F | BT | 10, 100 | 58, 74 |
| s-IL-6 | BT | 10, 100 | 53, 72 |
| sTNFα | BT | 10, 100 | 49, 66 |
| B cell proliferation | BT | 100 | 31 |
| MCP-1 | HDFSAG | 100 | 33 |
| IL-8 | HDFSAG | 10, 100 | 25, 44 |
| MMP-1 | HDFSAG | 100 | 23 |
| sIL-10 | HDFSAG | 10, 100 | 73, 81 |
| sIL-17A | HDFSAG | 10, 100 | 57, 74 |
| sIL-17F | HDFSAG | 1, 10, 100 | 46, 86, 83 |
| sIL-2 | HDFSAG | 10, 100 | 47, 68 |
| sIL-6 | HDFSAG | 1, 10, 100 | 33, 91, 96 |
| sTNFα | HDFSAG | 100 | 35 |
| sVEGF | HDFSAG | 1, 10, 100 | 90, 84, 60 |
| MCP-1 | /TH2 | 10, 100 | 21, 25 |
| sIL-17 | /TH2 | 100 | 23 |

Abbreviations: BCR, B-cell receptor; BT System, co-culture of CD19+ B cells and PBMC that utilizes BCR stimulation and sub-mitogenic TCR stimulation; HDFSAG system, co-culture of human primary dermal fibroblasts and PBMC that is stimulated with sub-mitogenic TCR levels; IL, interleukin; PBMC, Peripheral Blood Mononuclear Cell; SAG system, co-culture of endothelial cells and PBMC stimulated with mitogenic levels of TCR ligands; TCR, T-cell receptor; /TH2 system, co-culture of endothelial cells and Th2 blasts stimulated with TCR ligands and cytokines; Th2, T helper type 2; VEGF, vascular endothelial growth factor

In the BT co-cell culture system, which models chronic inflammatory conditions driven by B cell activation and antibody production, incubation of cells with 10 nM PTC299 resulted in a significant reduction in the levels of soluble (s)IgG, sIL-17A, sIL-17F, sIL-6, and sTNFα released from the cells after 72 hours of stimulation (range, 49% to 68%) (all p values <0.01) (Figure 4 and Table 2). The production of sIgG was also significantly decreased by 97% following 144 hours of stimulation (p <0.01). Similar to results obtained in the SAg co-cell culture system, 100 nM PTC299 significantly inhibited proliferation (31%) compared with control (p<0.05).

In the HDFSAg co-cell culture system, which models chronic T cell activation and inflammation responses within a tissue setting, PTC299 was associated with significantly lower levels of secreted MCP-1, IL-8, MMP-1, sIL-10, sIL-17A, sIL-17F, sIL-2, sIL-6, sTNFα and sVEGF compared with control (all p values <0.01) (Figure 4 and Table 2). PTC299 inhibition of cytokine production was particularly apparent for sIL-17F, sIL-6, and sVEGF, where a significant reduction in the levels of these cytokines was observed at as low as 1 nM PTC299 (range 33% to 90%; all p values <0.01). No significant differences between co-cultures treated with PTC299 or the control were seen with regard to proliferation or cytotoxicity.

In the /TH2 co-cell culture system, a model of mixed vascular Th1 and Th2 type inflammation that creates a pro-angiogenic environment promoting vascular permeability and recruitment of mast cells, basophils, eosinophils and T and B cells, significant inhibition of MCP-1 and sIL-17A production was seen as relative to controls (all p values <0.01) (Figure 4 and Table 2). Significant inhibition of MCP-1 production of 21% was observed when cells were incubated with 10 nM PTC299 (p<0.01). Similar to results obtained with the HDFSAg co-culture system, treatment with PTC299 did not result in a significant decrease in cell proliferation or increased cytotoxicity in the /TH2 system.

### PTC299 Inhibits IL-17 Production in Th17 cells

IL-17A and IL-17F are cytokines produced by activated T-helper-17 (Th17) cells that mobilize and activate neutrophils. In COVID-19, disease severity is positively correlated with levels of IL-17A, and the levels of this cytokine are increased in patients in the intensive care unit (Megna et al., 2020; Pacha et al., 2020). Results from the BioSeek analysis described above using the BT and HDFSAg co-cell culture systems demonstrated that PTC299 significantly inhibits the production of IL-17A and IL-17F. The ability of PTC299 to decrease IL-17A and IL-17F levels was further verified by assessing the effects of PTC299 on the production of IL-17A and IL-17F by Th17 cells, a subset of CD4 T cells that produce these cytokines (Martinez et al., 2012).

The ability of PTC299 to inhibit the production of secreted IL-1**7**A and IL-17F was assessed in a model system in which PBMCs were stimulated with human T-activator CD3/CD28 Dynabeads in a culture containing cytokines and antibody to induce Th17 differentiation. The model culture system was incubated with increasing levels of PTC299. Consistent with the BioMap results, PTC299 inhibited the production of IL-17A and IL-17F in tissue culture medium in a dose-dependent manner, with >70% inhibition being observed at doses of ≥75 nM (Figures 5A and 5B). The inhibition was specific for PTC299, as the inactive *R*-enantiomer of this compound, PTC-371 (Cao et al., 2019) and teriflunomide, a DHODH inhibitor with a relatively high EC_50_ (26.1 μM) (Xiong et al., 2020), did not inhibit the production of IL-17A and IL-17F. A different DHODH inhibitor, brequinar, also inhibited IL-17A and IL-17F production, but at substantially higher concentrations than PTC299. Addition of 100 μM uridine rescued the PTC299-dependent inhibition consistent with PTC299 acting on DHODH (Figures 5C and 5D).

**Figure 5.**
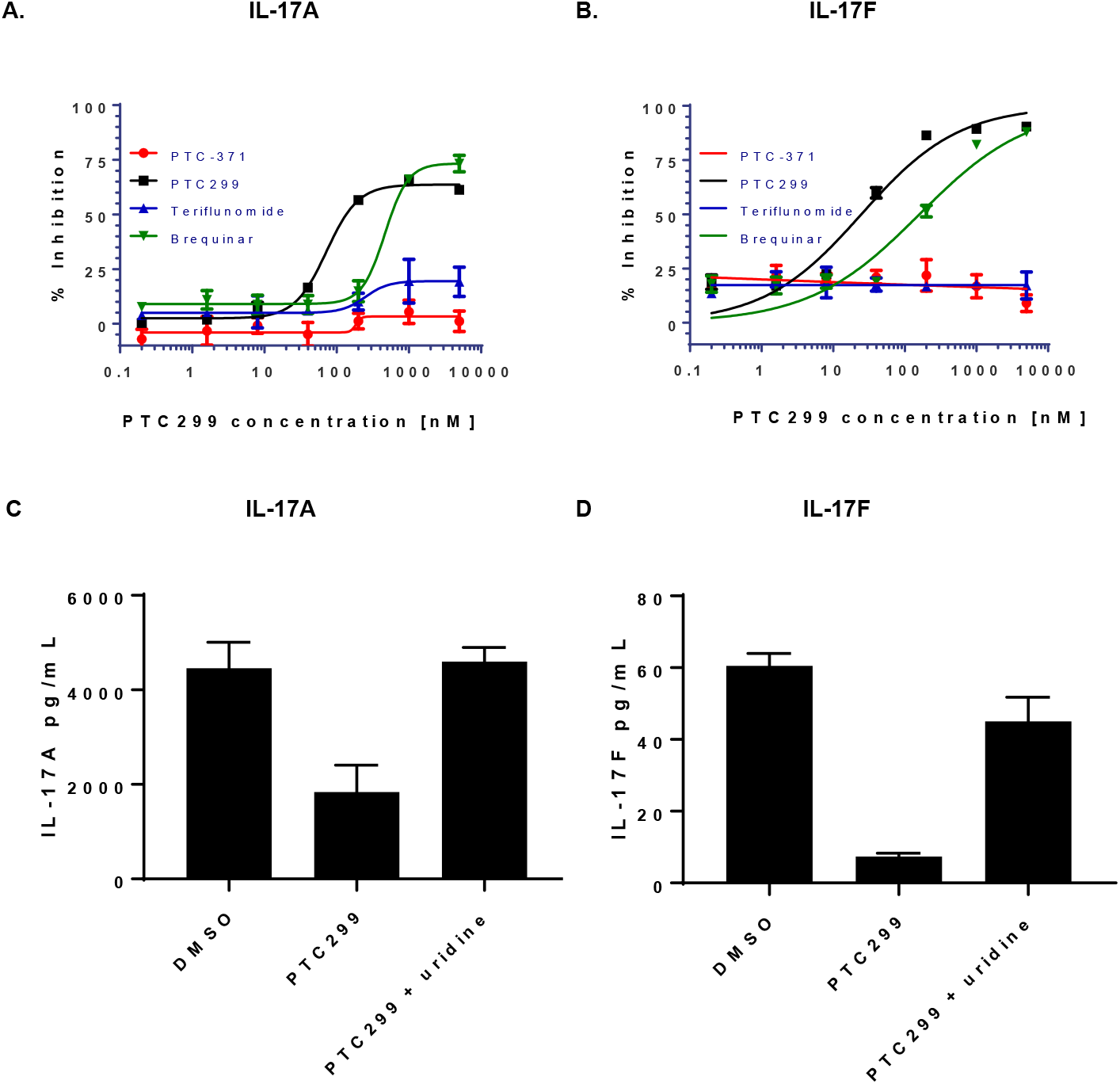
PTC299 Inhibits IL-17A and IL-17F Production in Th17 cells. Legend: (A-D) PMBCs were stimulated with T-cell activator CD3/CD28 Dynabeads and a combination of cytokines and antibodies to promote T-cell differentiation while blocking Th1 and Th2 differentiation. Cells were incubated with increasing concentrations of PTC299 and levels of (A) IL-17A and (B) IL-17F in the medium were measured by ELISA. Following PMBC stimulation and incubation in the presence of 1 μM PTC299 and100 μM uridine, levels of (C) IL-17A and (D) IL-17F in the medium were measured by ELISA. DMSO was used as a negative control Secreted IL-17F was reported as both pg/mL in the medium and as (pg/mL) normalized for cell number. Bars indicate standard deviation. Abbreviations: IL, interleukin; PBMC, peripheral blood mononuclear cell.

## DISCUSSION

COVID-19 is characterized by an early stage of viral replication followed in some cases by overproduction of inflammatory cytokines. Both the viral replication and the cytokine response are dependent upon pyrimidine nucleotides produced by the de novo biosynthesis pathway (Cheung et al., 2017; Xiong et al., 2020). The results presented here demonstrate that PTC299 has a dual mechanism of action that inhibits viral replication and attenuates the production of a subset of inflammatory cytokines. PTC299 potently inhibited SARS-CoV-2 replication in vitro, with an EC_50_ of 2.6 nM (31.6 nM based on the logarithmic curve), and an antiviral selectivity of >3,800. Inhibition of SARS-CoV-2 replication by PTC299 was prevented by the addition of exogenous uridine, which obviated the requirement for the de novo pyrimidine nucleotide synthesis pathway, consistent with the drug functioning as a DHODH inhibitor and with the fact that many RNA viruses require high concentrations of pyrimidine nucleotides to transcribe or replicate the viral genomes (Xiong et al., 2020). Accordingly, PTC299 also demonstrated broad-spectrum antiviral activity (Table 1).

A unique aspect of a DHODH inhibitor such as PTC299 is its ability to affect both viral replication and attenuate the cytokine storm (Breedveld and Dayer, 2000; Li et al., 2013; Xiong et al., 2020). The cytokine response associated with COVID-19 dramatically complicates the course of the infection in some patients and can result in ARDS with a high mortality rate. Experience with other viral infections indicates that treating the excessive inflammatory immune events downstream of the infection is paramount to patient recovery (Quartuccio et al., 2020). Studies presented here demonstrated that PTC299 inhibited production of multiple cytokines, including MCP-1, IL-6, TNFα, VEGF, and IL-17.

Inhibition of IL-6, IL-17, VEGF, and IgG by PTC299 may be of particular importance for treating COVID-19. IL-6 appears to play a key role in excessive cytokine production resulting from viral infection and pulmonary complications in COVID-19 (Ruan et al., 2020; Russell et al., 2020). Preliminary results from two small studies suggest inhibition of the IL-6 pathways by tocilizumab and siltuximab resulted in treatment benefit for COVID-19 patients (Gritti et al., 2020; Xu et al., 2020). Comparable to what was found for MERS-CoV and SAR-CoV infections, IL-17A has also been associated with disease severity and lung injury in COVID-19 (Pacha et al., 2020). Similarly, increased VEGF levels promote vascular permeability and leakage, helping in the pathophysiology of hypotension and pulmonary dysfunction (Teuwen et al., 2020). The attenuation of expression of these cytokines by PTC299 that result from SARS-CoV-2 infection may provide important and unique therapeutic benefits. The reduction in the levels of IgG production by PTC299 may also be a clinically meaningful aspect of PTC299 treatment; elevated IgG levels has been shown to be associated with the severity of COVID-19 (Tan et al., 2020; Zhao et al., 2020).

The novel dual mechanism of action of PTC299 distinguishes it from most other therapeutics being investigated in the clinic for the treatment of COVID-19, as many of these target either viral-specific processes or the immune response, but not both. Due to its dual mechanism of action, PTC299 is expected to be effective in treating both early and later stages of COVID-19. This contrasts with direct-acting antivirals that typically show their greatest effectiveness primarily in the early phase of the disease. Further, since PTC299 targets a cellular gene product (DHODH), and it is unlikely that viruses can overcome the need for pyrimidine nucleotides, the likelihood that the therapeutic effect of PTC299 will be compromised by the development of viral resistance is low. This may be particularly important for RNA viruses whose high mutation frequency often promotes evasion of direct-acting antivirals.

PTC299 is a highly potent inhibitor of SARS-CoV-2 replication, with an EC_50_ of 2.6 nM. Several other DHODH inhibitors have also shown activity against SARS-CoV-2 (Xiong et al., 2020), including teriflunomide, an FDA approved treatment for relapsing forms of multiple sclerosis. Teriflunomide has an EC_50_ of 26.1 μM, and similar to PTC299, has also been shown to have broad spectrum antiviral activity (Xiong et al., 2020).

Several other compounds that are under evaluation in the clinic for the treatment of SARS-CoV-2 infection have EC_50_ values reported to be the micromolar range. While these compounds have different mechanisms of action than PTC299, it is notable that the reported EC_50_ of remdesivir is 770 nM, that of chloroquine is 1.1 to 5.5 μM, and that of hydroxycholorquine is 720 nM (Choy et al., 2020; Sanders et al., 2020; Wang et al., 2020b; Xiong et al., 2020; Yao et al., 2020).

The key to a successful COVID-19 treatment is to not only have a potent molecule, but to also have a dose that can be delivered safely and that will sustain exposure in the blood to inhibit viral replication or infection. PTC299 is currently being evaluated to treat COVID-19 in the phase 2/3 study PTC299-VIR-015-COV19 (referred to as FITE19) (https://clinicaltrials.gov/ct2/show/NCT04439071?term=PTC299+AND+COVID&draw=2&rank=1). The dose being used is based on a PK/pharmacodynamic relationship obtained in monkeys and in cancer patients with neurofibromatosis type 2 (NF2), and the well-characterized PK profile from healthy volunteer and oncology studies (Cao et al., 2019)(data on file). The PTC299 dose in the FITE19 study is predicted to yield a C_ave_ of 1371 ng/mL and will thus be ~1100-fold higher than the drug’s EC_50_ and ~55-fold greater than the EC_90_ values against SARS-CoV-2. The PTC299 levels would also be ~12-fold higher than the 250 nM concentration that resulted in the 3-log reduction in the titer of SARS-CoV-2 in cultured Vero cells.

In summary, PTC299 is a highly potent inhibitor of SARS-CoV-2 replication that also suppresses production of a subset of pro-inflammatory cytokines, suggesting it has the potential to act through this dual mechanism to treat the viral and immune components of COVID-19. Due to its ability to block viral replication and cytokine production, PTC299 may be effective in treating both early and later stages of the disease. This contrasts with direct-acting antivirals that typically show their greatest effectiveness primarily in the early phase of the disease (McNicholl and McNicholl, 2001; Xiong et al., 2020) and with anti-inflammatory drugs that treat only the later phase of COVID-19 (Moore and June, 2020; Quartuccio et al., 2020). It has been argued that it is important to combine antiviral therapy with immune suppressants as using only compounds that modify the immune response to treat the cytokine storms may make viral clearance more difficult (Quartuccio et al., 2020). Importantly, PTC299 is orally bioavailable, has been extensively evaluated in human subjects, has well established PK and safety profiles and is generally well tolerated. These findings and prior clinical experience with PTC299 support the further development of this novel molecule for the treatment of COVID-19.

## ACKNOWLEDGMENTS

The authors would like to thank Leonid Yurkovetskiy, Mitchell White, and Baylee Heiden for thoughtful discussion and constructive advice and help with design and execution of SARS-CoV-2 experiments.

The study was supported by NIH grants R37AI147868 and R01AI148784 to J.L, a grant from the Evergrande COVID-19 Response Fund Award from the Massachusetts Consortium on Pathogen Readiness to J.L., and a grant from the Evergrande COVID-19 Response Fund Award from the Massachusetts Consortium on Pathogen Readiness to E.M. S.B. was supported by the Defense Threat Reduction Agency, Joint Science Technology Office, and S.P. and R.S were supported through Sponsored Research Agreement from PTC Therapeutics to University of Texas Medical Branch.

## AUTHOR CONTRIBUTIONS

Experimental design, methodology, and execution: J.L, S.P., E.M., R.S. S.B. C.S-D-C., E.L.S., Y.W, V.S., J.D.G., L.C., M.W., C.T., K.C-L.

Writing, review, and editing: J.L., S.P., E.M., R.S., S.B., J.D.G., M.W., C.T., N.A.N., J.M.C., M.P., E.M.W., K.OK, R.K., E.G. A.J., S.P.

Project oversight: N.A.N, J.M.C, S.P.

## METHODS

### Experimental Materials

#### KEY RESOURCES TABLE

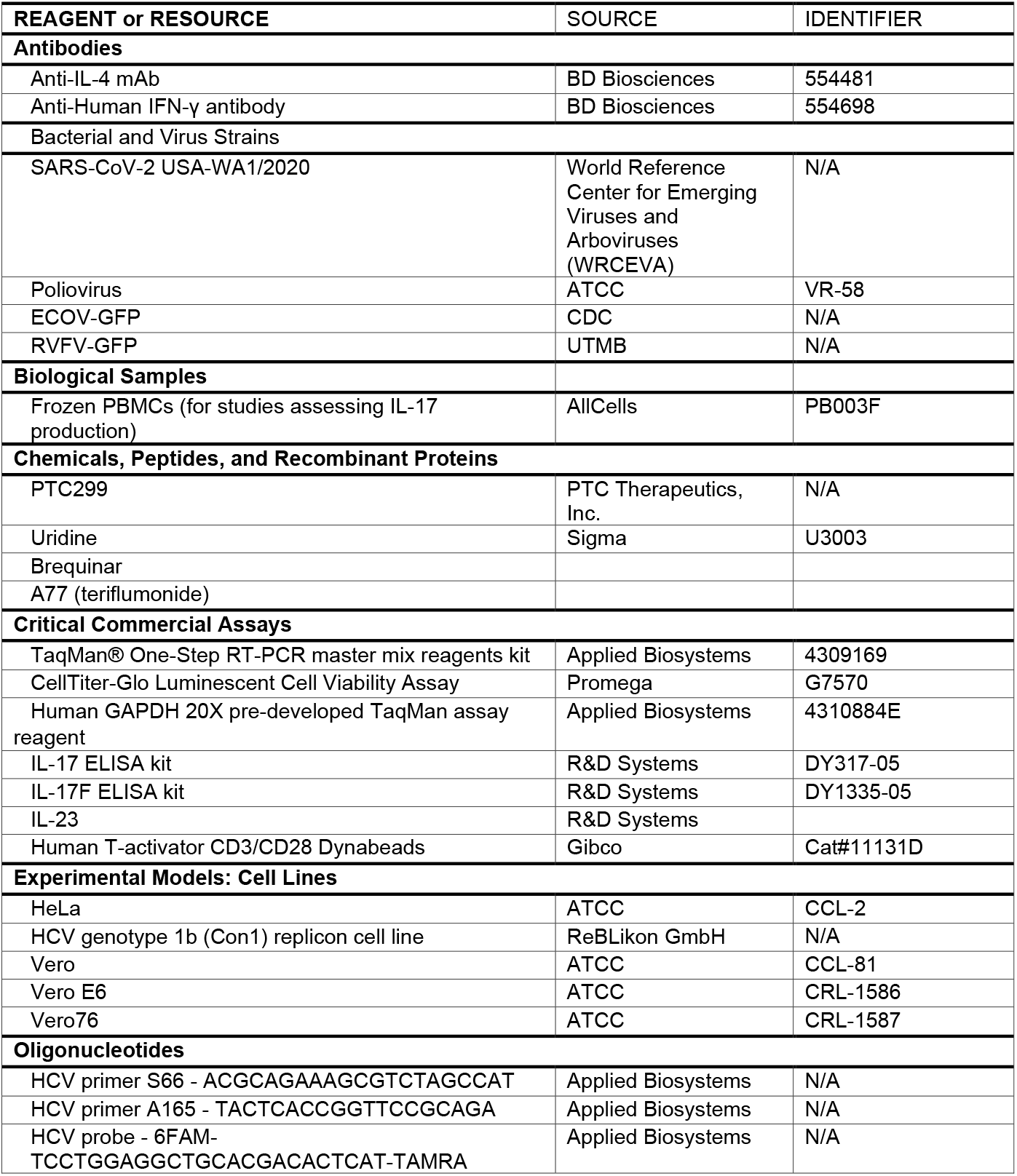
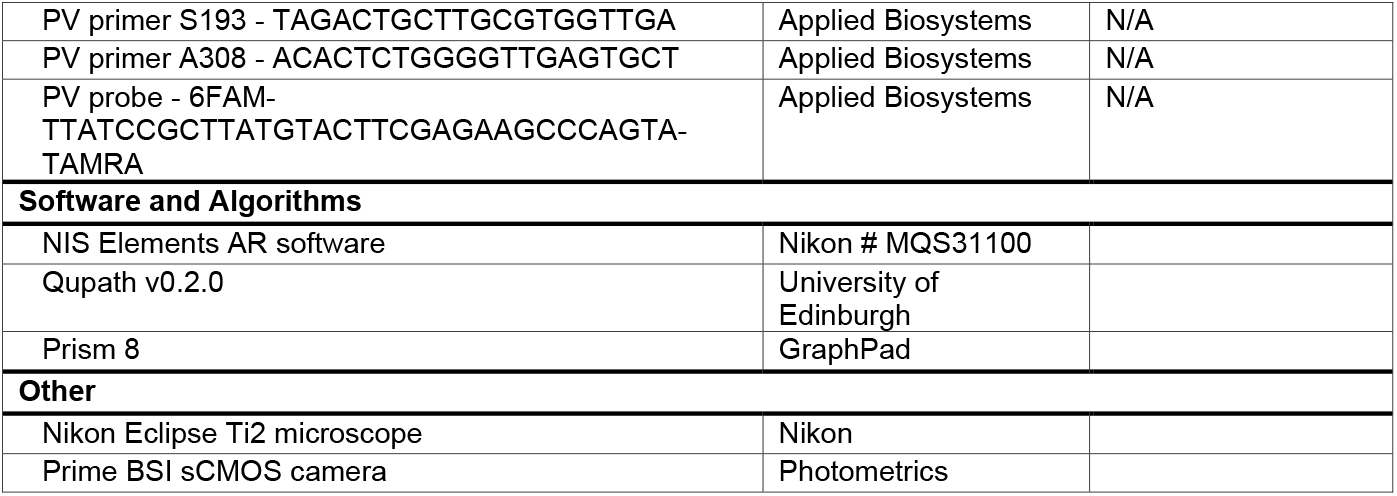

### Cell cultures

Human and primate cells HeLa (ATCC, CCL-2), Vero (ATCC, CCL-81), Vero E6 cells (ATCC, CRL-1586) were maintained at 37°C and 5% CO2 in Dulbecco’s Modified Eagle Medium (DMEM, Hyclone or GIBCO) supplemented with 10% Fetal Bovine Serum (FBS, Corning or GIBCO). Huh7 cell line harboring a genotype 1b replicon was maintained under similar conditions but with the addition of 0.25 mg/ml G418 (GIBCO). G418 was for replicon cell maintenance only and was removed during compound testing.

### Viruses

SARS-CoV-2 USA-WA1/2020 was provided by the World Reference Center for Emerging Viruses and Arboviruses (WRCEVA) and was originally obtained from the Centers for Disease Control and Prevention (CDC). The virus was subsequently titrated and passaged once more on Vero cells, grown in DMEM supplemented with 5% FBS and 1% penicillin and streptomycin solution. All experiments were conducted at the University of Texas Medical Branch (UTMB) approved biosafety level-three (BSL-3) laboratories and all personnel undergo routine medical surveillance.

Poliovirus (CCL-2) was obtained from ATCC. The received virus was amplified in HeLa cells, grown in DMEM supplemented with 5% FBS and 1% penicillin and streptomycin solution and the titer evaluated by plaque assay on HeLa cell monolayers.

### Detailed Methods

#### SARS-CoV-2 viral yield assay

Vero cells were pretreated with increasing concentrations (1 nM to 1 μM) of PTC299 or PTC299 with 100 μM uridine. After incubation at 37°C overnight, medium was removed, and cells were inoculated with SARS-CoV-2 at a multiplicity of infection (MOI) of 0.05. After 1 hour of incubation at 37°C inCO_2_, wells were washed 3 times with dilution medium and 1 mL of medium containing the indicated compound doses was added back to each well. Cells were incubated at 37°C and samples collected at 0, 16, and 24 hours post-infection. Collected timepoint medium was replaced with an equal amount of 1x compound or dilution medium. Samples were stored at −80°C until the day of analysis. The SARS-CoV-2 titer in Vero cells via 50% tissue culture infectious dose assay (TCID_50_) was performed for each sample collected at 0, 16, and 24 hours post-infection.

For cytotoxicity determination, Vero cells were plated at 1e5 or 2e5 cells/ml DMEM containing 5% FBS and 1% penicillin and streptomycin solution. Cells were treated with PTC299 at concentrations from 1 nM to 10 μM. At 16, 24, or 48 hours post-treatment, cytotoxicity was measured by Cell Titer-Glo Luminescent Cell Viability Assay (Promega, Madison, WI, USA).

#### Quantification of SARS-CoV-2 infection levels by quantitative immunofluorescence analysis

African Green Monkey Vero E6 cells were plated in 96-well plates at 1×10^4^ cells per well. The next day, cells were either pretreated with DMSO alone, increasing concentrations of PTC299 in DMSO or they were untreated for 30 minutes at 37°C and 5% CO2. Cells were subsequently infected with SARS-CoV-2 at a multiplicity of infection (MOI) of 0.1 by adding 10 μL of the diluted virus to each well. Experiments were conducted in triplicates using three wells of a 96-well plate per condition.

Two days post infection, cells were fixed in 10% neutral buffered formalin for at least 6 hours at 4°C and removed from the BSL-4 laboratory. The cells were permeabilized with 1:1 (vol:vol) acetone-methanol solution for 5 min at −20°C, incubated in 0.1 M glycine for 10 min at room temperature, and subsequently incubated in blocking reagent (2% bovine serum albumin, 0.2% Tween 20, 3% glycerin, and 0.05% NaN3 in PBS) for 20 minutes at room temperature. After each step, the cells were washed three times in PBS, and then incubated for 1 hour at room temperature with a rabbit antibody directed against the SARS-CoV nucleoprotein (Rockland Immunochemicals, Gilbertsville, PA). This antibody cross-reacts with the SARS-CoV-2 nucleoprotein. The cells were washed four times in PBS and incubated with goat anti-rabbit antibody conjugated with AlexaFluor488 (Invitrogen) for 1 hour at room temperature. 4’,6-diamidino-2-phenylindole (DAPI) (Sigma-Aldrich, St Louis, MO) was used at 200 ng/mL for nuclei staining.

Cells were stored at 4°C in PBS prior to imaging. Images were acquired using a with Photometrics Prime BSI sCMOS camera attached to a Nikon Eclipse Ti2 microscope, equipped with a Nikon N Plan Apo λ 4X/0.20 ∞/- WD20. Acquisition was driven using the NIS Elements AR software (Nikon). Four tiles were acquired per well and tiles were stitched together using the NIS Element AR Software. Two channels were acquired per tile using the Nikon C-FL DAPI and the Nikon C-FL GFP HC HISN Zero Shift filter sets, respectively.

Images corresponding to 4 tiles and 2 channels each, were imported into QuPath software v0.2.0 (https://qupath.github.io/) and subjected to quantitative image analysis using a custom-made script based on the Multiplexed Analysis procedure (https://qupath.readthedocs.io/en/latest/docs/tutorials/multiplex_analysis.html). SARS-CoV-2 positive cells were counted and their normalized percentage with respect to the total number of identified cells was plotted against the Log10 transform of the concentration of PTC299. The non-linear regression of the inverted sigmoid curve was calculated using GraphPad Prism 8.

Data management was performed using the OMERO.web client (v5.4.6-ice36-b87) and the OMERO server v5.4.6 (https://www.openmicroscopy.org/omero/.

#### PTC299 activity against RNA virus panel

For poliovirus antiviral assay, HeLa cells were seeded at 5000/well in 96-well plates the day before infection in DMEM containing 10% fetal bovine serum (FBS). Cells were treated with serially diluted test compounds 16 hours pre-infection. The final concentration of dimethyl sulfoxide (DMSO) in the media was 0.5%. Cells were infected with 0.1 plaque forming units per cell of poliovirus (PV) and incubated at 37°C. Twenty hours post-infection, cells were lysed, and quantitative reverse transcription-polymerase chain reaction (RT-qPCR) was performed using primers to the poliovirus genome and to human glyceraldehyde 3-phosphate dehydrogenase (GAPDH), a cellular reference gene, Predeveloped TaqMan^®^ assay reagents (Applied Biosystems, Foster City, CA). Concentration required for 50% inhibition of viral genomic RNA compared to mock treated control (EC_50_) was reported as the concentration of test compound required to cause a 50% reduction in PV RNA and was normalized to GAPDH levels in the same well. The concentration required for 50% inhibition of cellular GAPDH RNA compared to mock treated control (CC_50_) was reported as the concentration of test compound that caused a 50% reduction in GAPDH mRNA. The experiment was repeated 3 times for this study and average EC_50_ and CC_50_ values are reported.

HCV replicon studies were performed as previously described. Huh-7 (hepatocarcinoma) cells harboring the subgenomic HCV genotype 1b (Con1) replicon (referred to as 1b replicon cells) were obtained from ReBLikon GmbH (Gau-Odernheim, Germany). Replicon cells were plated at a density of 5000 cells per well in 96-well plates in DMEM containing 10% FBS. Serial dilutions of test compounds in DMSO were added to wells and cells were incubated at 37°C and 5% CO2 for 3 days prior to determining the extent of inhibition of HCV replicon RNA replication and the effect of the compound on cell proliferation. The final concentration of DMSO in the medium was 0.5%. Inhibition of replicon RNA replication was quantified using real-time RT-qPCR assay. At the end of compound treatment, the cells were lysed with Cell-to-cDNA lysis buffer (Ambion, Austin, TX). RT-PCR was carried out using TaqMan^®^ One-Step RT-PCR master mix reagents kit (Applied Biosystems) with HCV primers (sense S66 [ACGCAGAAAGCGTCTAGCCAT] and anti-sense A165 [TACTCACCGGTTCCGCAGA]) and probe (5’-6FAM-TCCTGGAGGCTGCACGACACTCAT - 3’ TAMRA) at a concentration of 100 μM. The effect of the compound on the abundance of the housekeeping gene GAPDH mRNA as a control for cytotoxicity was determined by using Predeveloped TaqMan^®^ assay reagents (Applied Biosystems, Foster City, CA). Both HCV replicon and GAPDH RNAs were amplified in the same well using ABI 7900HT (Applied Biosystems). The quantity of HCV replicon or GAPDH RNA from each sample was determined by applying PCR cycle-time values to a standard curve. The percent reduction of HCV RNA in the presence of test compound was calculated using the formula: [1-(compound treatment sample - background control) / (non-compound treatment control - background control)] X 100. EC_50_ and EC_90_ (50% and 90% effective concentrations, respectively) values were calculated by non-linear regression using Prism software (GraphPad, San Diego, CA). The test was repeated 6 times for this study and average EC_50_ and CC_50_ values are reported.

Evaluation of the effect of PTC299 on Ebola and Rift Valley Fever (RVFV) virus replication was performed at USAMRIID (Fort Detrick, MD). Assays were performed in 96 well plates and analyzed by high content screening. Test compounds were administered at a concentration range of 0.0032 to 2 μM in duplicate. Antiviral assays used a virus engineered to express green fluorescent protein (EBOV-GFP), and antiviral activity was determined by counting the number of GFP expressing cells compared to a mock treated control. Cytotoxicity was determined by counting the number of viable cells compared to mock treated control. For evaluation of efficacy against Ebola virus, Vero76 cells were seeded at 30,000 cells per well 1 day prior to infection and pretreated for 18 or 2 hours with test compound. Cells were infected at a multiplicity of infection of 5 with Ebola-GFP virus and incubated 48 hours at 37°C. For evaluation of efficacy against Rift Valley fever virus, HeLa cells were seeded at 10,000 cells per well 1 day prior to infection and pretreated for 2 hours with test compound. Cells were infected at a multiplicity of infection of 0.1 with RVFV-GFP virus and incubated 48 hours at 37°C.

#### Evaluation of inhibition of cytokine production

The activity of PT299 was evaluated in complex primary culture human cell systems in the BioMap assays profiling platform (from BioSeek, now part of Eurofins). Detailed protocols for the BioMap primary human cell cultures and coculture systems have previously been published (Berg et al., 2010; Berg et al., 2013; Melton et al., 2013; Shah et al., 2017). These systems consist of complex co-cultures of human peripheral blood mononuclear cells (PBMCs) pooled from healthy donors with early passage human primary fibroblasts, B-cells, or venular endothelial cells and are stimulated to recapitulate relevant signaling networks that naturally occur in human tissue or during specific disease states. In this study, four different co-culture systems were utilized. PTC299 was prepared in DMSO (final concentration ≤ 0.1%) and was added at the specified concentrations 1-hr before stimulation and remain in culture for 24-hrs (sAg system), 48-hrs (HDFSAg system), 72-hrs (BT system, soluble cytokines), or144-hrs (BT system, secreted IgG)).

##### SAg system

The SAg system is a co-culture of human umbilical vein endothelial cells (HUVEC) and peripheral blood mononuclear cells (PBMC) stimulated with mitogenic levels of T cell receptor (TCR) ligands to drive the polyclonal T cell stimulation (defined as 1X TCR ligand strength), expansion and elevated cytokine release that is characteristic of acute T cell immune activation and inflammation conditions. HUVEC cells were stimulated with cytokines (interleukin [IL]-1β, 1 ng/mL; TNF-α, 5 ng/mL; and IFN-γ, 100 ng/mL) and the superantigen Staphylococcal enterotoxin F from Staphylococcus aureus [Sag (20 ng/mL)]. The SAg system also captures the paracrine signaling between activated immune cells and inflamed endothelial cells that is relevant for the vascular inflammation.

##### BT system

The BT system is a co-culture of CD19+B cells and PBMC that utilizes B cell receptor (BCR) stimulation and sub-mitogenic TCR stimulation (defined as 1/1000th TCR ligand strength) to capture the T cell dependent B cell activation and class switching that occurs in germinal centers. The BT system models diseases and chronic inflammatory conditions driven by B cell activation and antibody production.

##### HDFSAg system

The HDFSAg system is a co-culture of human primary dermal fibroblasts and PBMC that is stimulated with sub-mitogenic TCR levels (defined as 1/1000th TCR ligand strength) to model chronic T cell activation and inflammation responses within a tissue setting.

##### /TH2 system

The /TH2 system is a co-culture of HUVEC cells and Th2 blasts stimulated with TCR ligands (1/10th ligand strength) and cytokines to model mixed vascular T helper type 1 (Th1) and Th2 type inflammation and create a pro-angiogenic environment that promotes vascular permeability and recruitment of mast cells, basophils, eosinophils and T and B cells.

For all 4 co-cell culture systems, PTC299 was added to cells at a concentration of 0.1, 1.0, 10, and 100 nM 1 hr prior to stimulation of the cells and was present during the 48-hr stimulation period. The effect of PTC299 on the levels of various biomarkers including cytokines, growth factors, expression of surface molecules, and cell proliferation were monitored. The levels of protein biomarkers were measured by ELISA. Cytotoxicity was quantified by Alamar blue reduction, which measures cell viability by evaluating the level of oxidation during respiration. Cell viability was quantified using Sulforhodamine B (SRB) assay. SRB bind cellular protein components and measure the total biomass and proliferation was measured by counting cells.

#### IL-17A and IL-17F ELISA

PBMCs (Allcells, Inc.) were stimulated with T cell activator CD3/CD28 Dynabeads, that are coated with anti-CD3 and anti-CD28 antibodies (Dynabeads; Gibco, Cat#11131D), in the presence of cytokines and antibodies to promote Th17 differentiation while blocking Th1 and Th2 differentiation. The final concentration of cytokines was as follows: IL-6 (10 ng/mL), IL-1β (10 ng/mL), TGF-β1 (5–10 ng/mL), IL-23 (10 ng/mL), and the neutralizing antibodies anti-IL-4 (10 μg/mL) and anti-IFN-γ (10 μg/mL) in culture medium.

For all experiments, a total of 40 μL of cells (1 x 105 cells) were added per well in a 96-well tissue culture plate. Subsequently, 10 μL of washed Dynabeads or medium were added to each well and the cells were incubated at 37°C, 5% CO_2_ for 3 to 4 hours. Following incubation,, 25 μL of 4x cytokine/Ab mixture (described above) was as added to each well (for the “no stimulating control”, 25 μL medium was as added) and 25 μL of PTC299, PTC-371 (the inactive enantiomer of PTC299) (Cao et al., 2019), brequinar, A77, or medium was added. Cells were incubated at 37°C, 5% CO2 for a total 88 hours. After 88 hours, the medium was collected and analyzed by an enzyme-linked immunosorbent assay (ELISA) kit (R&D System: IL17 Cat#DY317-05 and IL17F Cat#DY1335-05) according to manufacturer’s instruction.

#### Statistical Analysis

Data was analyzed using GraphPad PRISM.

