## Supplemental Figure 1 for "The DHODH Inhibitor PTC299 Arrests SARS-CoV-2 Replication and Suppresses Induction of Inflammatory Cytokines"

### SUPPLEMENTAL INFORMATION

**Supplemental Figure 1. PTC299 inhibition of SARS-CoV-2 is via inhibition of DHODH**

**A. 18 hours preincubation with PTC299 prior to viral infection**

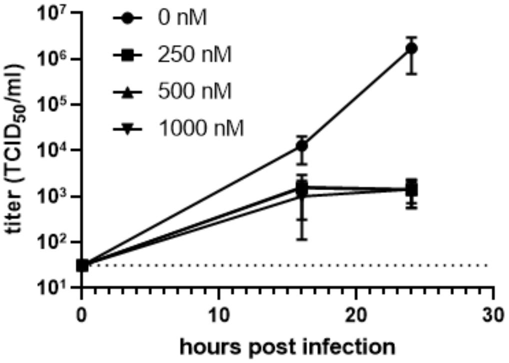


**B. Addition of PTC299 at same time as viral infection**

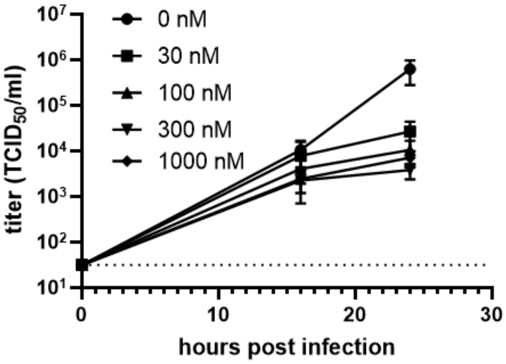


**C. Treatment with PTC299 2 hours after viral infection**

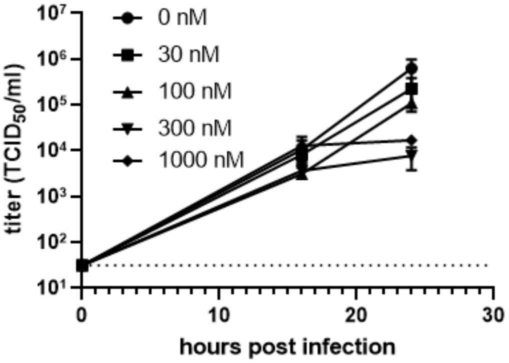


Legend: Vero cells were treated with PTC299 at the indicated concentrations for 18 hours before, at the same time as, or 2 hours after infection. Cells were inoculated with SARS-CoV-2 (USA-WA1/2020) at a MOI of 0.05. Viral titer was determined by collecting the medium from the infected cells and assessing the concentration required for 50% infectivity of Vero cells (the 50% tissue culture infectious dose assay or $TCID_{50}$). Data is plotted as the mean and standard deviation of 3 independent replicate.
